# Does Food Insecurity exist among Farm Households? Evidence from Ghana

**DOI:** 10.1101/2021.02.04.429712

**Authors:** Patricia Pinamang Acheampong, Elvis Agyei Obeng, Monica Opoku, Lydia Brobbey, Bernard Sakyiamah

## Abstract

Household food security exists when households have physical, social and economic access to sufficient, safe and nutritious food at all times that meets their dietary needs and food preferences for an active and healthy life. Food security remains a serious challenge for many households in Ghana and the situation is even more prevalent among smallholder farmers. Using data collected from 2,603 farm households across Ghana and employing an ordered probit model the determinants of food security among farm households were assessed. The food security indicator-Food Consumption Score (FCS) which combines diet diversity, frequency of consumption and relative nutritional importance of different food groups was used for the analysis. Results indicated that farm households (76%) across Ghana were within the acceptable household food consumption groups. Nonetheless, 19% and 6% of farm households respectively were within the borderline and poor food consumption groups respectively. Further analysis revealed the determinants of food security to include experience, gender, improved variety adoption, access to credit and location. The suggestion is that government and private institutions should create an enabling environment to enhancing production capacities, economic and social resilience to improve on food security and nutrition.

## Introduction

[1] defines Food security as “a situation in which all people at all times have physical and economic access to sufficient, safe and nutritious food which meets their dietary needs and food preferences for an active and healthy life’’. This definition integrates stability, access to food, availability of nutritionally adequate food, biological utilization of food as well as food system stability. In order to achieve food security and food system stability, farm households cultivate more than one crop on a piece of land. Farm diversity has a more direct effect on household dietary diversity in subsistence farming than households that produce mainly for income [2]. The quality of diet consumed by farm households is dependent for example on whether the household is producing high income-generating crops or livestock. [3] opined that diversity in crops production is significantly related to households’ dietary diversity and is more associated with households’ consumption from own-produced food rather than consumption of market- purchased food. According to [4], most of the undernourished and poor people are smallholder farmers, so the move to further diversify production on these smallholder farms is an expedient tactic to enhance dietary quality and diversity. Consequently, there is a need to diversify agricultural production so that a wide variety of food is available and accessible to poor segments of the population [5].

Food insecurity is mostly caused by insufficient calorie intake and malnutrition due to poor quality diet and low intake of essential nutrients such as vitamins and minerals [6]. These nutrients cannot be obtained in a single diet. Hence dietary diversification is critical. Dietary diversity according to [7] is the increase in the variety of foods within and across food groups to ensure sufficient intake of vital nutrients to promote good health. Household diets which contain more food groups are more likely to provide sufficient nutrients the body requires daily. Availability, accessibility and utilization which are the aspects of food security are linked to dietary diversity [7, 8]. [9] noted that higher rates of food insecurity and malnutrition are experienced by rural smallholder farmers in comparison to urban households. Farm production diversity is associated with greater household dietary diversity [2].

Aside farm production, several other factors influence the food security status of farming households.[10] found farm production diversity to be positively associated with household’s dietary diversity in their study on agricultural diversity and dietary diversity in Eastern India. Again in India, [11] observed that dietary diversity of farmers’ households was enhanced by better education; and higher crop diversity Other income-generating pathways like the sale of more crops to gain income for the purchase of more diverse food from the markets are region-dependent contributors to farmer household dietary diversity though vary from region to region.[4] in their study of determinants of dietary diversity and nutrition in Tanzania posited that household dietary diversity is significantly influenced by the wealth status of the households, household size, gender of the household head, spatial and temporal availability of diverse foods, diversity of agricultural production and agrobiodiversity, as well as personality and tradition.

In a study to find the determinants of household dietary practices in rural Tanzania [12]found that households’ dietary diversity is significantly influenced by the wealth status of the household, access to media, ownership of land, reduced risk of food insecurity and literacy. Household’s dietary diversity is also strongly influenced by farm production diversity according to [13]. [6] in their study of production diversity and dietary diversity in smallholder farm households suggested that the association between farm production diversity and dietary diversity was inherently confounded by market access. Additionally, [14] in their study of determinants of household food security in the Sekyere-Afram Plains District of Ghana found that farm size, off-farm income and credit access significantly influenced food security of farmers. In their study of food security in the Savannah Accelerated Development Authority Zone of Ghana, [15] found that food security could be determined by multiple crop production, yield and commercialization.

Although farm households are the main food suppliers, they are frequently the most food insecure, as they face biophysical and socioeconomic challenges [16]. The need to assess food security status of food crop producers and to suggest policy directions is imperative. Previous studies conducted on farm households in Ghana, such as[14] and [15] have all conducted case studies. This study used rich data collected on farm households across Ghana. The assessments of food security in this study provide broad information about status of food security and its determinants among farm households in Ghana. This study therefore had its objectives to evaluate food security using food consumption scores which combines diet diversity, frequency of consumption and relative nutritional importance of different food groups and also to assess factors that determine food security of food crops producers using an ordered probit approach.

## Materials and methods

### The study area^1^

Data for this study were collected from farm households across Ghana. All regions of the country were surveyed from August 2016 to February 2017. Ghana’s agriculture is predominantly on a smallholder basis where majority of farm holdings are less than 2 hectares in size. Annual average temperatures range from 26.1°C in places near the coast to 28.9°C in the extreme north. Day time temperatures may rise above 40°C in the north. There are two rainy seasons in the south from March to July and from September to October (bimodal rainfall system). The northern part of the country, on the other hand, has only one rainy season, from May to October (uni-modal rainfall system) [17].

### Sampling and data collection

A multi-stage sampling technique was employed to select regions, districts and villages. Regions and districts were purposively selected for the study. Afterwards, farm households were randomly selected from Enumeration Areas (EAs) within the selected villages. EAs were used for the proportional sampling of farm households. Enumeration areas (EAs) were obtained from the Ghana Statistical Service. Crops producing districts were collected from regional and district directorate of Ministry of Food and Agriculture. In each district, 4 EAs were selected and 7 farm households were contacted. A total of 99 districts and 2772 farm households were contacted across the country. However due to incomplete information 169 were discarded and data from 2603 households were utilized.

The selected households were interviewed face to face with structured questionnaires, which were carefully designed and pre-tested. In addition to farm and farmer data, data on various food items were collected. The household-level dietary section of the questionnaire was answered by the person responsible for food preparation in the household. Food consumption at the household level was captured through a 7-day recall.

### Measuring farm household food security

Food security is multidimensional and thus presents a variety of measurements. Several indicators have been developed as proxies for food security. A comprehensive all-encompassing measure of food security would be that measure that is valid and reliable, comparable over time and space, and which captures different elements of food security [18]. Nevertheless, the complexity of food security, as a crosscutting discipline, has engrossed the challenge to finding a summative (or ‘gold standard’) measure of household food insecurity [19]. [20] includes thirty- three indicators in its recommended list of indicators for measuring food insecurity access alone.[21] asserted that the need to finding a simple and realistic measure of household food insecurity that can be labelled as “golden rule” combining rigour and statistical efficiency to conclude food insecurity from the household level upwards is important. Some measures of food security from various literature [19,22] are as enumerated in Table 1. [24] describes the causes of household food insecurity vulnerability to be “difficult to measure”.

**Table 1.**
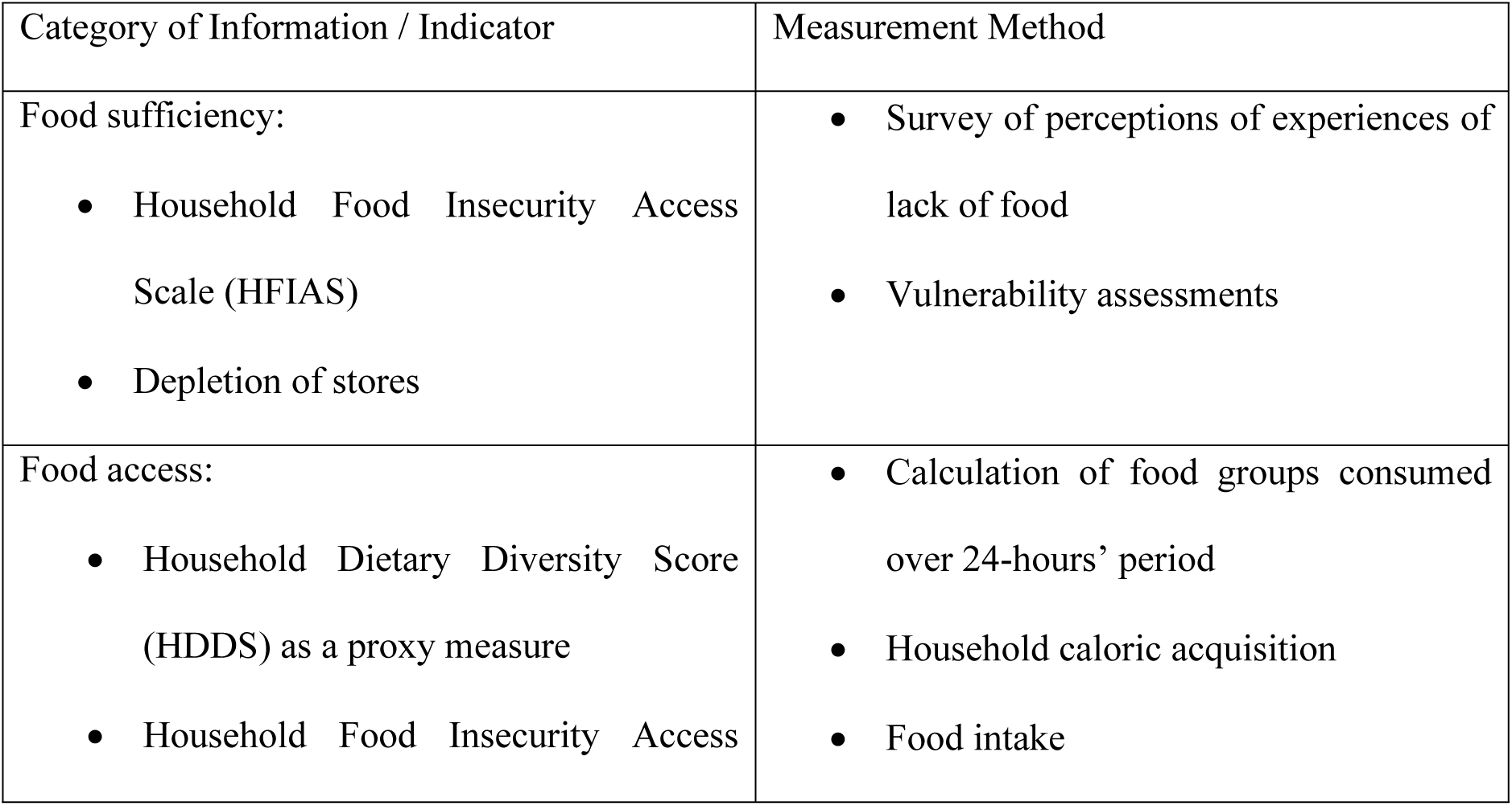
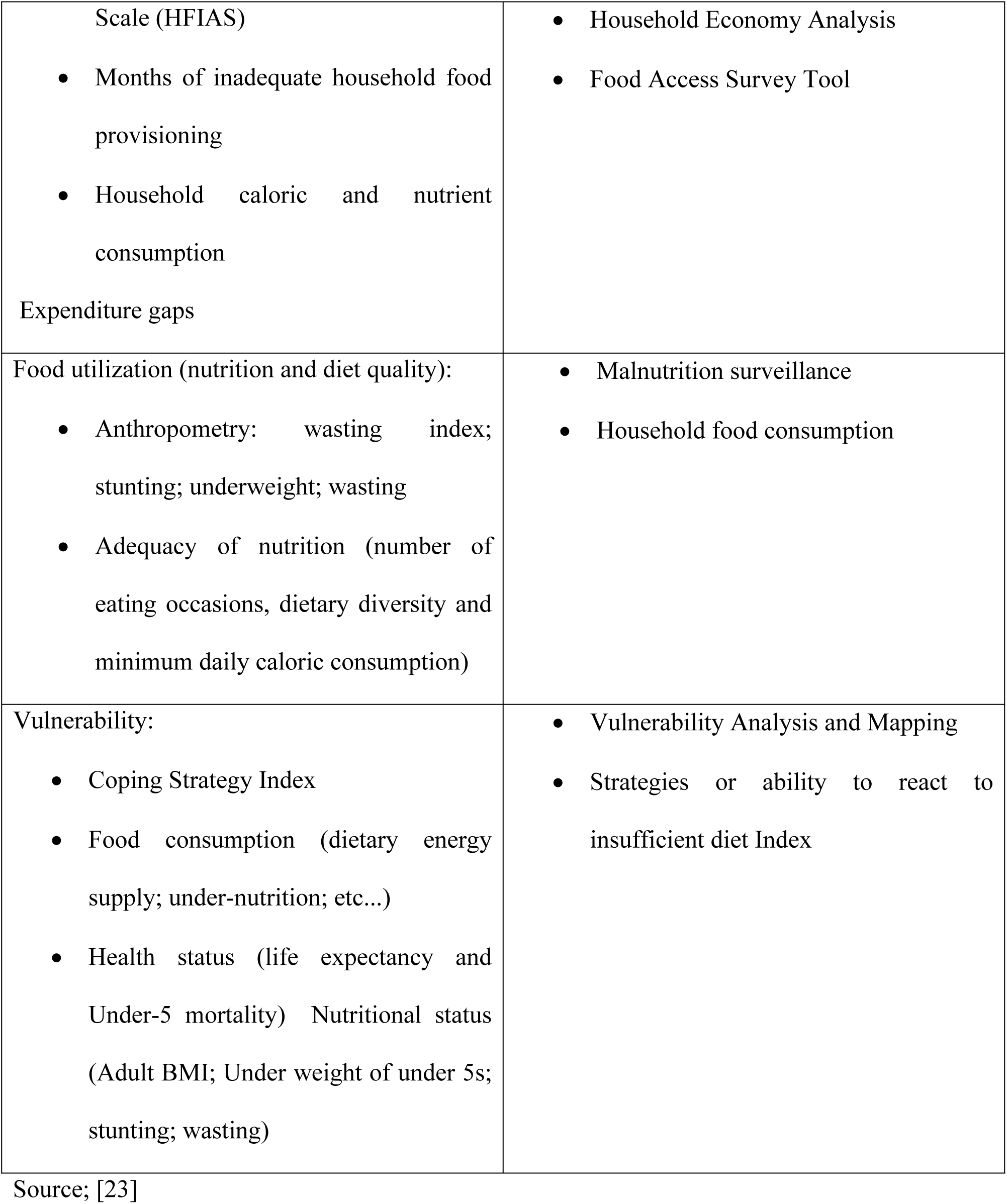
Measures and indicators of household food insecurity

It further categorizes the causes of food insecurity as: (a) availability of quantity and quality of household food; (b) physical and economic access to food. [25] adds the dimension of food utilization to the equation. These authors reinforce views by others stating that “adequate food availability at the aggregate level is a necessary, although not sufficient, condition to achieve adequate food access at the household level, which in turn, is necessary but not sufficient for adequate food utilization at the individual level”. [26] notes that food availability is influenced by stocks available locally, imports, food aid and local food production. Food production is influenced by household characteristics and other institutional factors. Food access is affected directly by food production, the market and cash transfers (i.e. government/private remittances or kinship support), and indirectly by food availability through food production prices [23].

This study used household food consumption score (FCS) as a measure of farm household food security. The FCS was developed by World Food Program (WFP) and it is a composite score based on dietary diversity, food frequency, and relative nutritional importance of different food groups [24]. Data were collected from the list of food items available in the country. Respondents were asked about frequency of consumption of specific food items (in days) over a recall period of the past 7 days. Food items were grouped into 8 standard food groups with a maximum value of 7 days/week [24]. The consumption frequency of each food group was multiplied by an assigned weight (Table 2) that is based on its nutrient content. Those values were then summed up obtaining the Food Consumption Score (FCS). FCS was then classified into three categories: poor consumption (FCS = 1 to 28); borderline (FCS = 28.5 to 42); and acceptable consumption (FCS = >42).

**Table 2.**
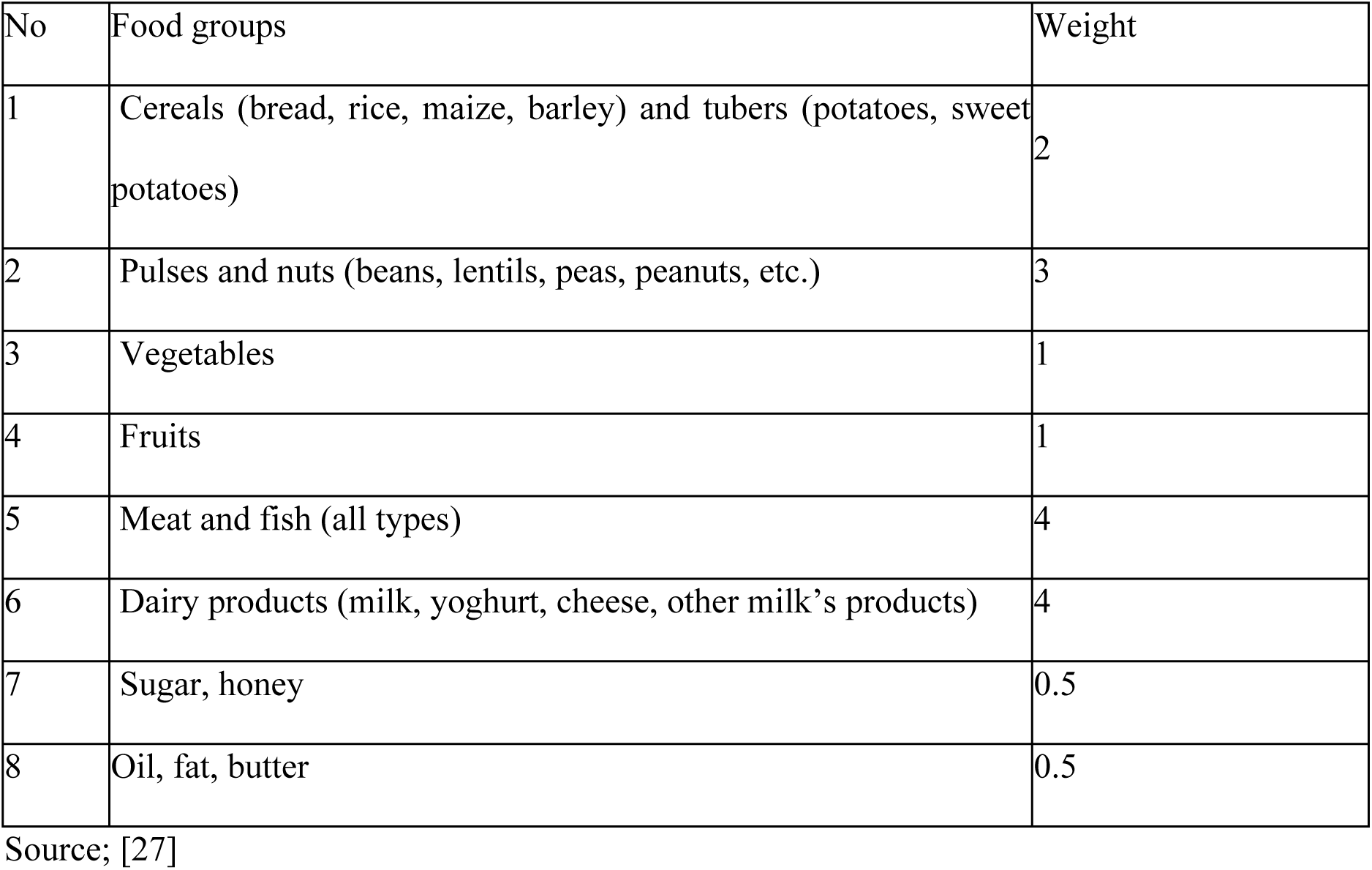
Food Items, Food Group and Weight

| No | Food groups | Weight |
| --- | --- | --- |
| 1 | Cereals (bread, rice, maize, barley) and tubers (potatoes, sweet potatoes) | 2 |
| 2 | Pulses and nuts (beans, lentils, peas, peanuts, etc.) | 3 |
| 3 | Vegetables | 1 |
| 4 | Fruits | 1 |
| 5 | Meat and fish (all types) | 4 |
| 6 | Dairy products (milk, yoghurt, cheese, other milk's products) | 4 |
| 7 | Sugar, honey | 0.5 |
| 8 | Oil, fat, butter | 0.5 |

Source; [27]

### Empirical model

An ordered probit model was used due to the ordinal nature of the dependent variable. The dependent variable was FCS which was categorical and had 0=poor; 1= borderline and 2=acceptable which indicated the food security status of farm households. [28] opined that granting that the outcome is discrete, the multinomial logit or probit models would fail to account for the ordinal nature of the dependent variable. Whereas the logit assumes a logistic distribution of the error term, the probit assumes a normal distribution [29]. According to [28] the logistic and normal distributions generally give similar results usually. Moreover, [30] point out that the ordered probit is the most used model for ordered response data in applied econometric work. The ordered probit is used in this study as a result. Following [29], an ordered probit model is written as:

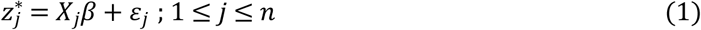

where *z*_*j*_ is a continuous, latent variable, *X*_*j*_ is a 1 × *k* vector of explanatory variables, *β* is a *k* × 1 vector of unknown parameters, and the *ε*_*j*_ are assumed to be independently and identically distributed with a probability density function (pdf) denoted *f*(*ε,θ*) with distributional parameters *θ*. The above model could be consistently estimated using OLS if *z*_*j*_ was observed and *E*⟨*ε*_*j*_│*X*_*j*_⟩ = 0,∀*j* [29]. Due to the discrete-choice nature of the data, OLS will result in heteroskedastic errors and predicted probabilities that may fall outside the range of (0, 1) for each outcome described. As a result of this problem, the maximum likelihood estimation is often used to estimate the unknown parameters. Consider the observed variable *z*_*j*_:

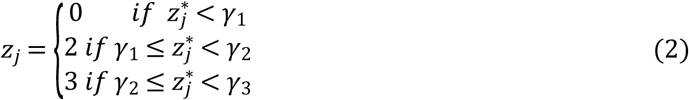

The probability of observing a particular outcome, for 1 ≤ *j* ≤ *i* is given by:

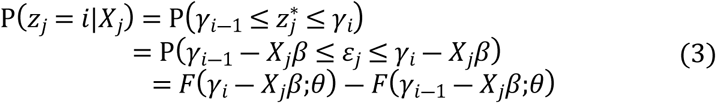

where F is the cumulative distribution function for *ε*_*j*_, *γ*_0_ = *γ*_*I*―1_ = −∞, and *γ*_*i*_ = ∞.The presence of F leads to a maximum likelihood estimation framework written as:

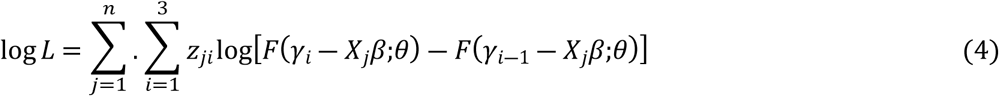

This log-likelihood function is maximized with respect to *β, θ* and the cut points *γ*_1_ < *γ*_2_. In the case of two discrete outcomes, the log-likelihood function in(4) simplifies to the binary choice model with one cut point which is normally set to be 0 to achieve identification of the intercept term [29].

[28], pointed out that since there is no meaningful conditional mean function and the marginal effects in the ordered probability models are not straightforward, the effects of changes in the explanatory variables on cell probabilities are normally considered. These are given by:

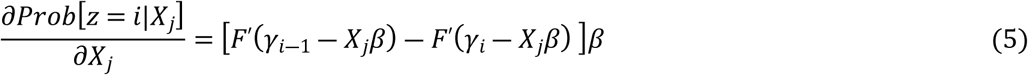

Consequently, the empirical model of this study is specified as:

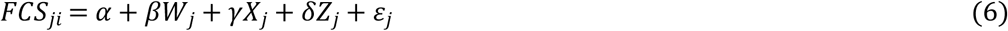

where FCS is food consumption score used as food security proxy; *j* represents a household, *i* (*I* = 0, 2, 3) represents the three categories of alternative dependent ordered variables indicating (i) whether a household falls within poor food consumption group category, (ii) whether a household falls within borderline food consumption category, and (iii) whether a household is within the acceptable food consumption category group. *W, X* and *Z* are, respectively, socioeconomic, food production, institutional and location characteristics hypothesized to influence food security (Table 3); *β*, *γ, α,δ* are parameters to be estimated.

**Table 3.** Descriptive statistics of respondents by Region

| Variables | AR<br>(N=34<br>5) | BA<br>(N=49<br>0) | CR<br>(N=28<br>1) | ER<br>(N=35<br>2) | GA<br>(N=8<br>4) | NR<br>(N=29<br>9) | UE<br>(N=19<br>6) | UW<br>(N=12<br>3) | VR<br>(N=35<br>1) | WR<br>(N=8<br>2) | Pooled<br>(N=26<br>03) |
| --- | --- | --- | --- | --- | --- | --- | --- | --- | --- | --- | --- |
| Gender |  |  |  |  |  |  |  |  |  |  |  |
| Male | 64.10 | 71.80 | 69.00 | 75.90 | 83.30 | 96.30 | 94.90 | 99.20 | 73.20 | 54.90 | 76.90 |
| Female | 35.90 | 28.20 | 31.00 | 24.10 | 16.70 | 3.70 | 5.10 | 0.80 | 26.80 | 45.10 | 23.10 |
| Marital status |  |  |  |  |  |  |  |  |  |  |  |
| Single | 6.10 | 7.10 | 5.70 | 6.20 | 4.80 | 2.70 | 8.70 | 3.30 | 4.80 | 7.30 | 5.80 |
| Married | 81.70 | 86.10 | 85.40 | 86.90 | 88.10 | 94.30 | 90.30 | 94.30 | 89.50 | 84.10 | 87.70 |
| Divorced | 5.80 | 3.70 | 4.60 | 3.10 | 4.80 | 1.70 | 0.50 | 1.60 | 3.10 | 3.70 | 3.40 |
| Widowed | 6.40 | 3.10 | 4.30 | 3.70 | 2.40 | 1.30 | 0.50 | 0.80 | 2.60 | 4.90 | 3.20 |
| Residence status |  |  |  |  |  |  |  |  |  |  |  |
| Native | 57.10 | 58.60 | 64.10 | 62.20 | 70.20 | 91.00 | 100.0<br>0 | 95.90 | 81.50 | 65.90 | 71.80 |
| Settler | 42.90 | 41.40 | 35.90 | 37.80 | 29.80 | 9.00 | 0.00 | 4.10 | 18.50 | 34.10 | 28.20 |
| Education Level |  |  |  |  |  |  |  |  |  |  |  |
| None | 27.80 | 35.10 | 29.50 | 20.20 | 27.40 | 76.60 | 57.70 | 65.90 | 41.60 | 50.00 | 40.50 |
| Up to JHS | 22.60 | 26.70 | 42.00 | 45.20 | 59.50 | 10.00 | 28.10 | 18.70 | 28.80 | 28.00 | 29.50 |
| Up to SHS | 44.30 | 35.30 | 26.30 | 31.80 | 8.30 | 12.70 | 12.20 | 12.20 | 27.40 | 20.70 | 27.20 |
| Tertiary<br>(excluding<br>university) | 3.50 | 2.00 | 1.40 | 2.30 | 4.80 | 0.70 | 1.50 | 2.40 | 1.70 | 1.20 | 2.00 |
| Tertiary<br>(University) | 1.70 | 0.80 | 0.70 | 0.60 | 0.00 | 0.00 | 0.50 | 0.80 | 0.60 | 0.00 | 0.70 |
| Access to<br>credit | 40.00 | 49.98 | 39.86 | 53.13 | 55.94 | 57.53 | 59.18 | 64.23 | 49.29 | 41.46 | 49.90 |
| Extension access | 77.10 | 81.02 | 77.58 | 78.41 | 86.90 | 82.61 | 81.63 | 78.05 | 77.49 | 65.85 | 79.10 |
| Age (mean) | 48.35 | 48.33 | 47.16 | 46.75 | 52.25 | 44.16 | 48.39 | 43.75 | 44.26 | 46.96 | 46.84 |
| Years of education (Mean) | 8.21 | 6.94 | 6.99 | 8.23 | 6.88 | 2.47 | 3.47 | 3.22 | 6.07 | 4.87 | 6.15 |
| Household size(mean) | 6.75 | 7.72 | 6.29 | 6.86 | 6.77 | 10.61 | 9.45 | 9.69 | 7.77 | 7.06 | 7.83 |
| Farming Experience(Mean) | 22.81 | 21.42 | 21.72 | 22.43 | 26.46 | 25.49 | 27.96 | 22.97 | 21.36 | 23.57 | 22.95 |

Note: AR is Ashanti Region; BA is Bono Ahafo region; CR is Central Region; ER is Eastern Region; GA is Greater Accra Region; NR is Northern Region; UE is Upper East region; UW is Upper West Region; VR is Volta Region; WR is Western Region

## Results and Discussion

### Descriptive statistics of data used in the regression analysis

Table 3 presents the descriptive statistics of data used in the analysis. Generally results showed that the respondents were dominated by males and they constituted about 77%. This was expected as most farm owners are males. The situation is worst in the three Northern regions of Ghana. The results revealed that Upper West, Northern and Upper East regions had as high as 99%, 96% and 95% of respondents respectively being males. In the three Northern Regions of Ghana, the customary laws do not permit women to own land [31], though both men and women undertake agricultural activities. Gender is anticipated to influence food security as male-headed households are more food secured than female-headed households [32]. Majority (88%) were married and were mostly natives from their respective locations. Marital status is expected to influence food security positively [33, 34]. On level of education, about 41% of the respondents had no formal education, about 30% had up to Junior High School (JHS)level of education and 27% had up to Senior High School(SHS) level of education.[35] found out that farm households in rural areas were mostly illiterates in their National census conducted on farm households in Ghana. Average age of a respondent was 47 years.

The average household size was about 8 persons, higher than national average of 5 [35]. The influence of household size on food security cannot be predetermined as large household size may add more pressure on household in terms of the number of people required to feed [36]. On the contrary, household size may mean the availability of family labour for other off farm activities that may increase incomes boosting food security situation of a household [37]. About 50% of the respondents had access to credit and 79% reported that they had access to extension. Access to institutional factors is very important to increasing production of farm produce and thus increasing food access. Credit may increase the probability of a household’s ability to procuring production inputs as seeds, chemicals, and hiring of labour [38], which could improve production and thus the household food situation. Access to credit by households was therefore predicted to positively correlate with household food security status.

### Farm Household’s dietary diversity

A Household Dietary Diversity (HDD) was estimated in order to seek understanding of the food groups consumed by farm households. The various food groups that different households consumed were used to compute the HDD in a 7-day recall period. Various food items of farm households were grouped into eight (8) standard food groups [24]. Fig.1 and Fig. 2 present farm household’s dietary diversity across Ghana. The results showed that cereals and root crops and vegetables were consumed every day by 96% and 90% respectively the seven day recall period. Ghanaian food system consists largely of cereals and root crops and local vegetables which are part of everyday culinary prepared in homes in both rural and urban areas. Vegetables were consumed everyday by about 91% of farm households. Data from the [17] noted that Ghanaians have the tradition of eating cereals and root and tuber crops. The results also showed that consumption of pulses (cowpea, groundnut, beans) and fruits were not as common as roots and tubers and cereals. However about 35% and 25% of the respondents consumed pulses and fruits respectively every day during the 7-day recall period. Per capita consumption of these crops is low in Ghana [17].

Fig 1. This is the Fig 1 Title. This is the Fig 1 legend.

**Fig 1.**
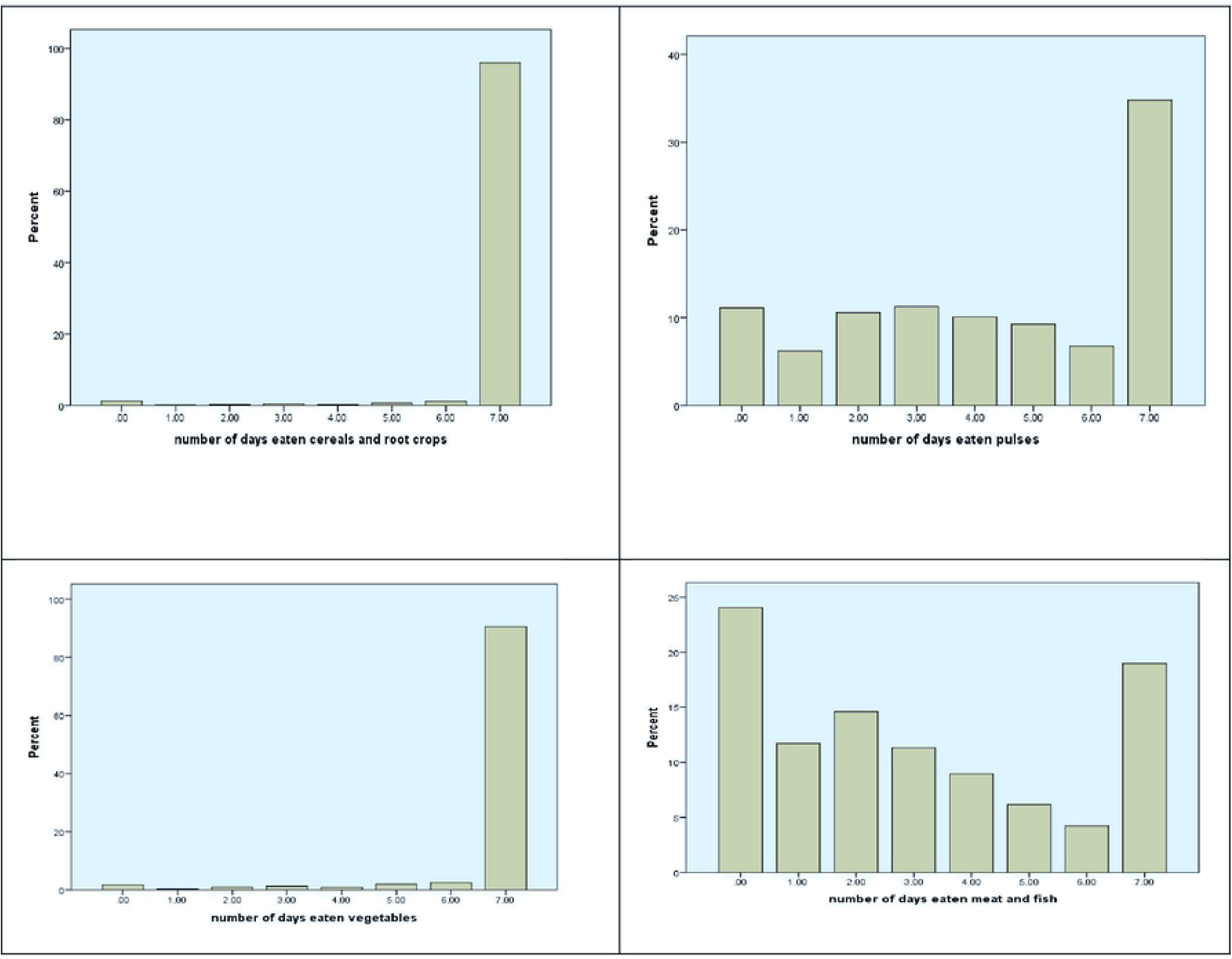
Consumption of Cereals and root crops, pulses, vegetable and meat and fish food groups

**Figure 2.**
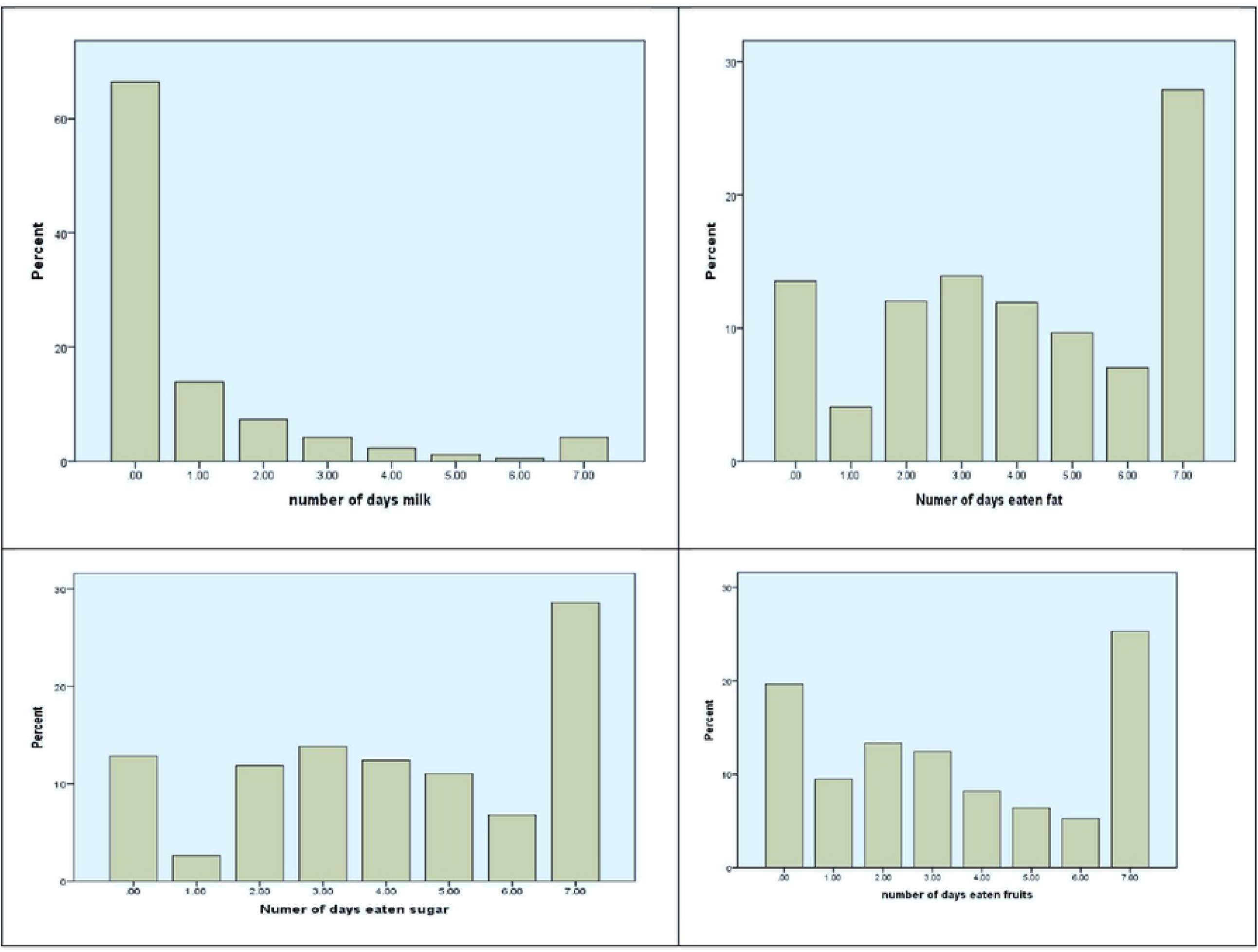
Consumption of Milk, fats, fruits, and sugars

Results from Fig. 2 revealed that most farm households in Ghana did not consume milk. About 66% of the respondents did not consume milk at all over the 7 day recall period and 24% had no meat or fish during the same period. Respondents also participated in fats and sugars to a certain level during the recall period. Nonetheless the results have revealed that if households did not produce a food item the frequency of consumption decreased. [39] found similar results in their study of Agricultural transformation and food and nutrition security in Ghana where households consumed own produce frequently compared with purchased food items.

Fig 2. This is the Fig 2 Title. This is the Fig 2 legend.

### Farm household’s Food Consumption score

The dietary diversity score does not show the amount (quantity) of food a household consumes. A Household Food Consumption Score (HFCS), a frequency-weighted HDDS, was further estimated as an indicator of dietary diversity and frequency of consumption by use of the frequency consumption of eight food groups previously mentioned. Fig 3 shows distribution of household food consumption score by regions. Results showed that majority (76%) of farm households across the country were within the acceptable HFCS.

**Fig 3.**
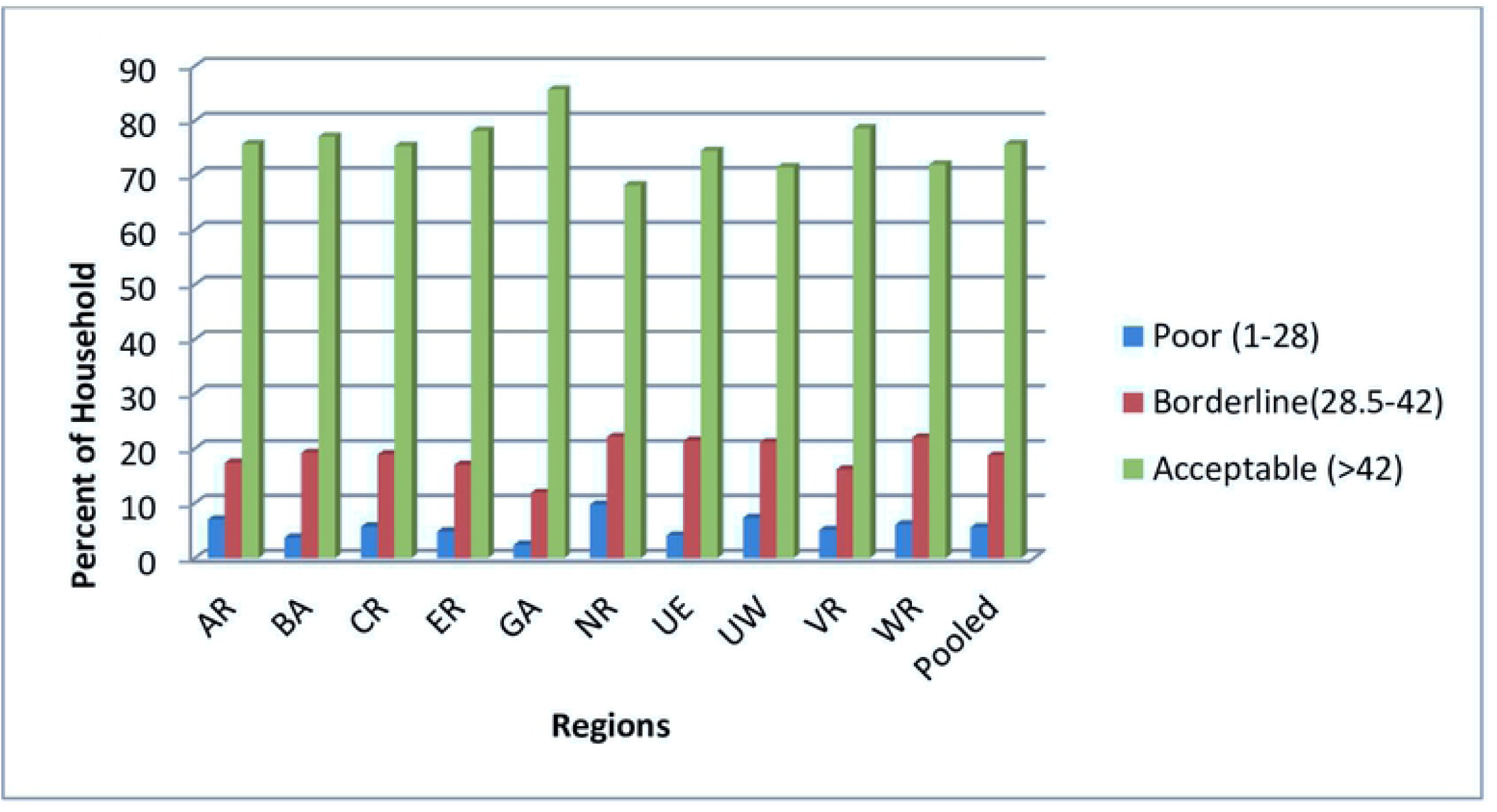
Distributions of Farm Household Food consumption score by region. Note: AR is Ashanti Region; BA is Bono Ahafo region; CR is Central Region; ER is Region; GA is Greater Accra Region; NR is Northern Region; UE is Upper East region; UW is Upper West Region; VR is Volta Region; WR is western Region

Fig 3. This is the Fig 3 Title. This is the Fig 3 legend.

Results also showed that majority of farm households in each of the regions had acceptable HFCS. Although there were substantial numbers (19%) of farm households on borderline HFCS across the regions, Northern, Western, Upper East and Upper West regions had 22.1%, 22%, 21.40% and 21.1% of members respectively on borderline HFCS higher than the rest of the regions. They are therefore moderately food insecure [27]. Northern, Upper East and Upper West regions have unfavorable climate in terms of rainfall and temperature. They have one season of rainfall and scorching temperatures which are not the case in the rest of the country. The implication is that some farmers from that part of Ghana may not have the luxury to produce diverse crops that contribute to food security. [27] stated that households that cultivate at least three different types of crops have better food consumption score than those that only cultivate one type in the Northern part of Ghana in their study of food security situation in the region. The Western region is known for the production of cash crops at the expense of food crops and thus the result was not surprising. We also discovered that there were some farm households across the country with poor HFCS, implying that they are severely food insecure. Generally about 6% of the respondents across the country reported severe food insecurity. [40] reported that about 5% of Ghana’s population was severely food insecure, a result which corroborates with results from this study.

### Determinants of food security among farm households

Table 4 presents results of determinants of food security among farm households in Ghana. The coefficients of the ordered probit estimates do not represent the magnitude of the effects of the explanatory variables, therefore the marginal effects are discussed. Results revealed that experience in farming decreased the probability of falling into the poor and borderline food consumption groups but increased the probability of falling into the acceptable food group. The possible explanation is that farmers with years of experience may have larger farm holdings and may also be practicing mixed farming thus making it possible to access all food groups with ease.[41] found a positive relationship between farming experience and food security status in their study of food security status among farm households in Nigeria. Our results showed that Up to JHS level of education decreased the probability of severe and moderate food insecurity of farm households but increased the likely of being food secured. Some level of education is important in ensuring diversity of food consumption in farm household. Similar result was obtained by [42] in their study of Household Food Security Status and Its Determinants in Maphumulo Local Municipality, South Africa where education positively influenced the food security status of households.

**Table 4.**
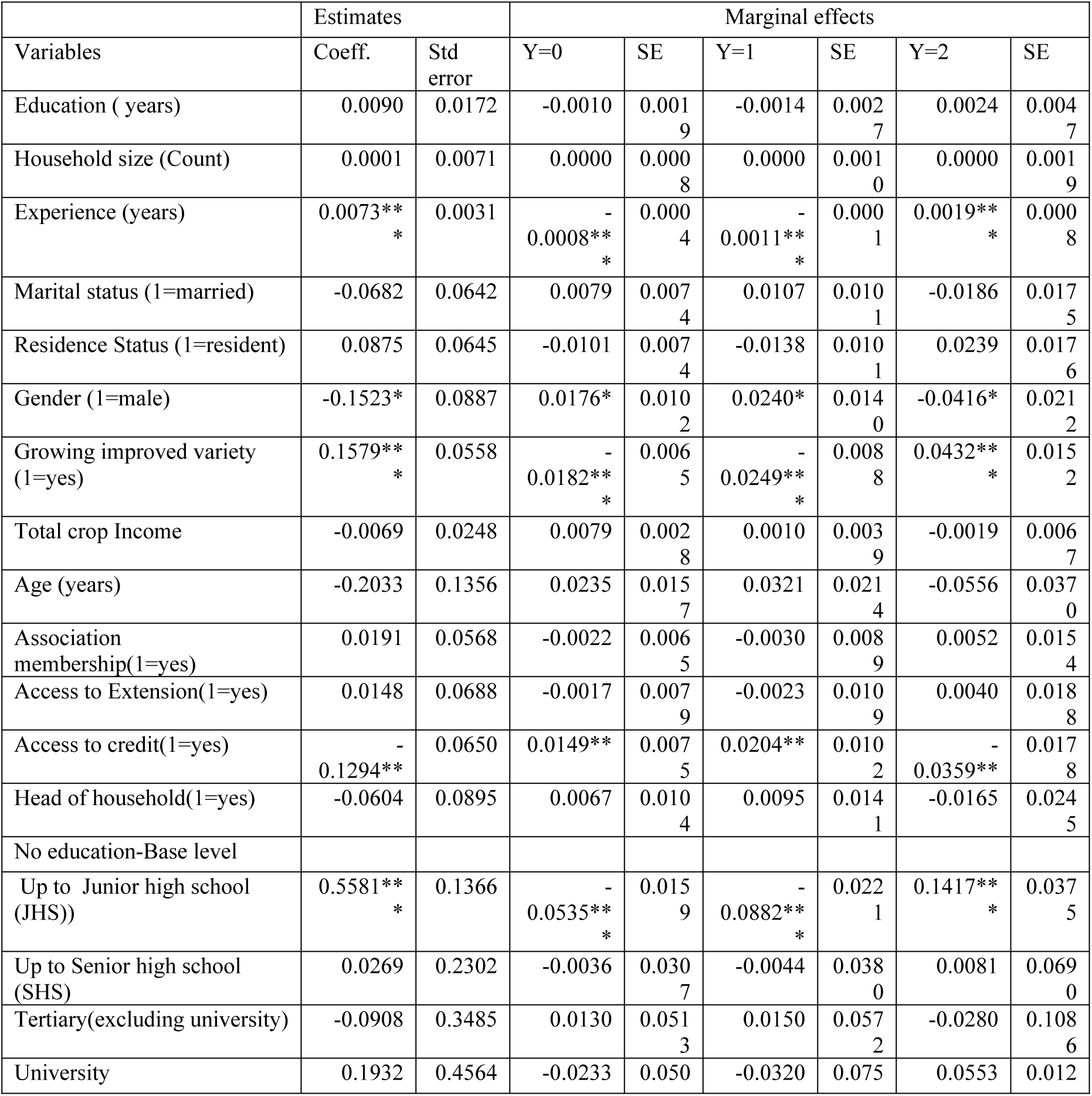
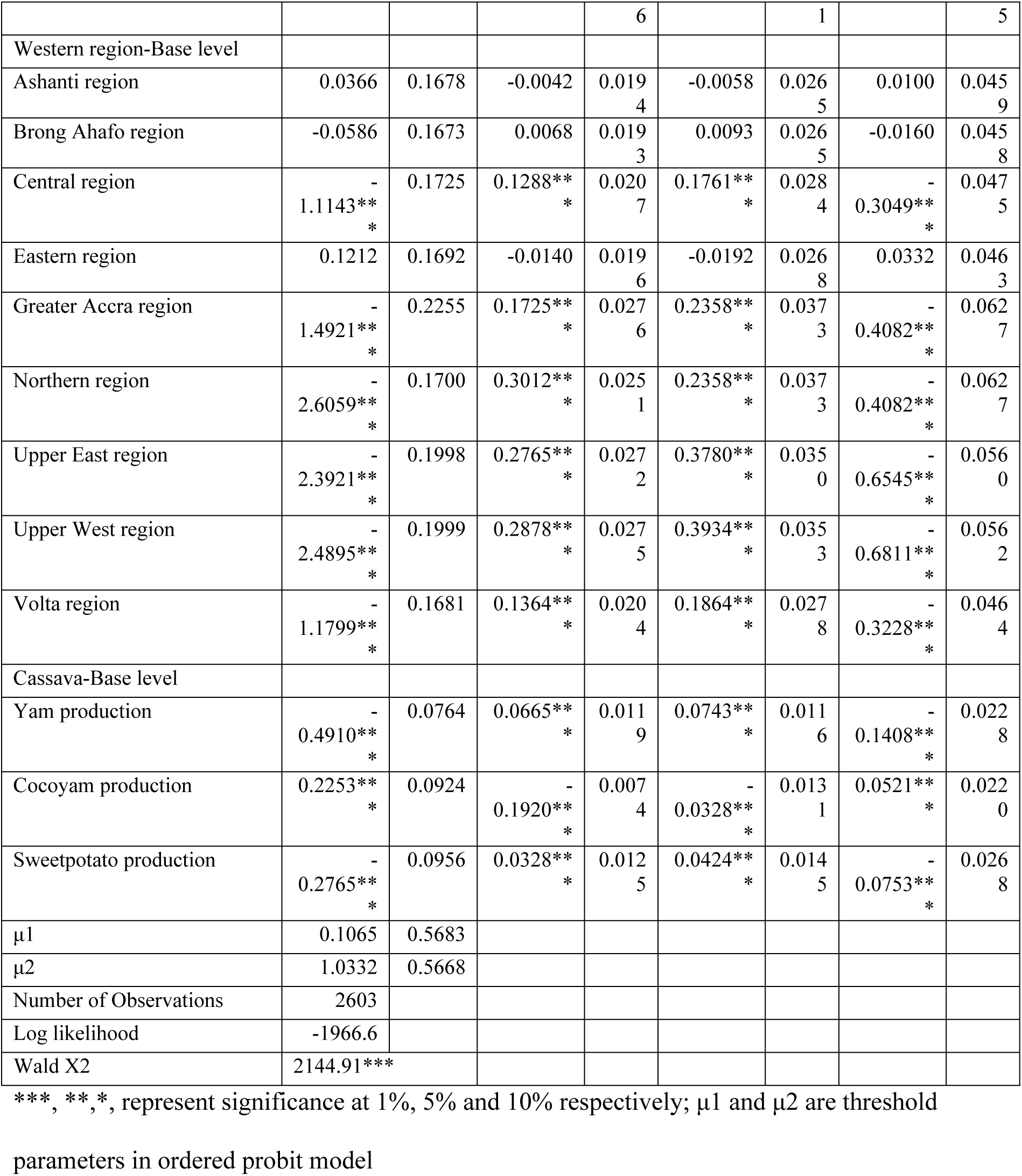
Ordered probit estimates of determinants of food security of farm households

We found that male headed farm households were more likely to be poorly food secured (poor food group) and moderately food secured (borderline food group). Nonetheless, female headed farm households were more likely to be highly food secured (acceptable food group). Female headed households mostly produced food crops that are mainly eaten by the household. They would only sell the surplus from production. The reverse is true for the male headed households who would sell most of their produce.

Our results discovered that growing of improved varieties decreased the probability of being severely food insecure and moderately food insecure but increases the probability of being highly food secure. Improved varieties are high yielding and those that produce them are likely to have enough for the household and also for sale to be able to purchase other food items. [43] found out in his study of Agrifood Systems and Sustainable Nutrition that adoption of modern crop varieties in increases yields and thus increases food availability and also improves agricultural profits and incomes of smallholder farmers. The results also revealed that production of roots and tuber crops

Access to credit was associated positively with farm household food insecurity, both severe and moderate but was negatively associated with acceptable food group. The suggestion is that farmers that have access to credit may use it for purposes other than farming. They may also not use credit for purchase of food items. [14] in their study of Determinants of Household Food Security in the Sekyere-Afram Plains District of Ghana found significant positive relationship of credit access on household food security. Their result is in contrast with the results of this study. However [44] found similar results on farm households in Northern Ghana where credit access significantly increased food insecurity of farm households. In Maphumulo Local Municipality, South Africa, [42] found credit access negatively influenced the food security status of households in their study of Household Food Security Status and Its Determinants, corroborating with this study result.

Farm households who are root crop producers are more likely to experience severe and moderate food insecurity and less likely to experience little or no food insecurity. Results showed that producers of yam and sweetpotato were likely to be severely and moderately food insecure than producers of cocoyam. This observation, though counter-intuitive, is pointing to whether conditions in those areas. Yam and sweetpotato producers are scattered in the transition and savannah agroecological zones where rainfall patterns are most erratic and thus may not able to produce enough for sale and for purchase of other food items. Cocoyam producers are mostly found in the forest agroecological zones where rainfall pattern is somehow reliable. They may thus be able to get enough harvest for sale and also capable of diversity food consumption.

As regards location, our results showed that farm household food security is associated with all regions except Ashanti, Bono Ahafo and Eastern regions. The results revealed that farm households in Central, Greater Accra, Northern, Western, Upper East, Upper West and Volta were more likely to be severely and moderately food insecure. This is not surprising as these regions fall within the forest transition and savannah agro-ecological zones where whether conditions are not always favorable. Farmers may not produce enough in those areas for consumption and for sale to purchase diverse food items.

## Conclusions and Implications

This study had its main goals to assessing farm household’s food security status and evaluating factors determining food security. We first assessed household dietary diversity to determine the number of individual foods consumed over 7-day period and then food consumption score as a proxy indicator of current food security of farm households. We also employed an ordered probit technique to evaluate factors influencing food security. Our results from household dietary diversity indicated that cereals and root tubers and vegetables food groups were the most consumed among farm households. Dairy products (milk) were the least consumed food group among farm households. Food security may be compromised as households perhaps are not meeting their micronutrient needs due to lack of inclusion of diary products. Farm households mostly consumed what they produce and the suggestion is that mixed farming, the practice of animal husbandry and crop production should be encouraged. As regarding household food consumption score (HFCS), the results revealed that 6% of the farm households across Ghana were within the poor HFCS. The implication is that food security remains an issue. Government agencies and non-governmental organizations should pay more attention to it by encouraging and educating households to do the needful to remain food secure.

The odds of being food secure by farm households were determined by factors that include, gender, experience, education level, and access to credit, improved variety cultivation and location. Our result showed that females were more food sure than males. The suggestion is that equitable access to inputs (credit access, labour acquisition, planting material, etc) and infrastructure be made available to all farmers to increase production and contribute to food security. Farming experience and some level of education (up to JHS level) determined food consumption score significantly. During dissemination of productivity enhancing technologies experience farmers could be targeted since they may be more receptive and open to trying new practice due to their ownership of vax lands and experience.

The general understanding is that education is important in uptake of improved technologies. Education determining food security suggests that educated members of farm households are able to diversify their livelihood portfolios, which could improve supply of diverse food items. Cultivation of improved variety increased the possibility of being food secure. Farmers’ ability to produce high yielding crop varieties increases home supply and increases sale of produce for purchase of other food options.

Credit access affected food consumption score negatively and significantly. The results showed that credit access reduced the probability of being food secure. Farm households may not be securing enough credit for farm production and may also not be securing credit for purchase of food items. Other purposes are assumed for accessing credit; therefore more education on credit management is important to give proper direction.

## Acknowledgements

We wish to acknowledge the funding provided by the World Bank through the West Africa Agricultural Productivity Programme for the study. Also the staff of CSIR-Crops Research Institute who assisted the data collection are greatly appreciated.

## Footnotes

1 The study was conducted when Ghana had 10 regions

